# Genes responsible for the formation of asymmetrical mandibular teeth in the giant mealworm beetle, *Zophobas atratus*

**DOI:** 10.1101/2025.01.10.632356

**Authors:** Kouhei Toga, Kakeru Yokoi, Toru Togawa

## Abstract

The mandibles of coleopteran insects are typical examples of external morphological asymmetry in animals. The developmental molecular mechanisms underlying the external morphological asymmetry remain to be elucidated compared to those of the internal asymmetric organ. In the tenebrionid beetle, *Zophobas atratus*, the inner teeth of adult mandibles are asymmetric, while the pupal mandibles are nearly symmetric. The asymmetric morphogenesis is assumed to be caused by the left-right regulatory differences of inner teeth formation, but even the developmental mechanisms involved in the inner teeth *per se* remains unknown. In this study, we investigated the morphogenetic process involved in the formation of inner teeth and searched for genes responsible for this process. Morphological observation showed that the inner teeth of adults formed during 0-5 days after pupal molting. Left mandibles possessed the characteristic second inner teeth, whereas the right mandibles exhibited invaginations between apical and second inner teeth. We performed RNAi for candidate genes and successfully identified four genes (*paired, alistaless, Lim1*, and *mlpt*) involved in the formation of inner teeth. Among them, *mlpt* RNAi had a strong effect on the morphology of the right mandibles, suggesting its involvement in the asymmetrical formation of mandibles.

## INTRODUCTION

Morphological asymmetry has independently occurred many times during the evolution of the animal kingdom (Palmer, 2009). The mechanisms underlying symmetry breaking in bilateral animals have been one of the main focuses of research in evolutionary developmental biology. Left-right asymmetry occurs in both external and internal organs. The molecular mechanisms involved in left-right asymmetry have mainly been investigated using internal organs of vertebrates (Coutelis et al., 2008; Grimes and Burdine, 2017). The findings revealed that twists of organs generate the asymmetric positioning of internal organs and that left-right asymmetry in gene expression executes symmetry breaking. *Nodal, Lefty-1*, and *Lefty-2* are representative genes expressed on the left side of vertebrate embryos (Tsukui et al., 1999; Shiratori and Hamada, 2014). The asymmetric positioning of internal organs has also been demonstrated in the fruit fly, *Drosophila melanogaster*, similar to vertebrates (Ligoxygakis et al., 2001; Hozumi et al., 2006; Spéder et al., 2006), even though a *Nodal* homolog was not found in the Arthropoda genome, including *D. melanogaster* (Bulm and Ott, 2018). It is also known that retinoic acid (RA) is involved in left-right asymmetry in vertebrates. In mice, RA controls the left-side expression of *Lefty-1* (Tsukui et al., 1999). In insects, juvenile hormone (JH), a retinoic-like hormone, plays a role in genitalia looping asymmetry (Ádám et al., 2003).

The asymmetry also occurs in the differences of organ size or morphology in a pair of external organs, as seen in the claws of lobsters. The molecular mechanisms underlying asymmetry in a pair of organs currently remain unclear. Insect mandibles are typical examples of the morphological asymmetry in a pair of organs (Okada et al, 2008; Toki and Togashi, 2011; Montealegre-Z et al., 2013; Sato et al., 2017). Although several genes involved in mandibular formation have been identified in the red flour beetle, *Tribolium castaneum* (Angelini et al., 2012; ibeetle base: Dönitz et al., 2015), and the stag beetle, *Cyclommatus metallifer* (Gotoh et al., 2017), the genes responsible for the formation of mandibular asymmetry remain unknown.

A tenebrionid beetle, *Zophobas atratus*, does not metamorphose under crowded conditions, whereas solitary conditions induce metamorphosis (Quennedey et al., 1995; Aribi et al., 1997; Quennedey and Quennedey, 1999; Toga et al 2021). Therefore, it is easy to manipulate the timing of pupation. Since the body size of this species is markedly larger than that of *T. castaneum*, a model beetle, morphological observations are also easy. Mandibles of *Z. atratus* adults show the asymmetric inner teeth (Fig. 1A-D). We aimed to reveal the genes associated with inner teeth formation in *Z. atratus*. In this study, we investigated the potential role of candidate genes that were predicted to be involved in asymmetric inner tooth formation in *Z. atratus*.

**Figure 1.**
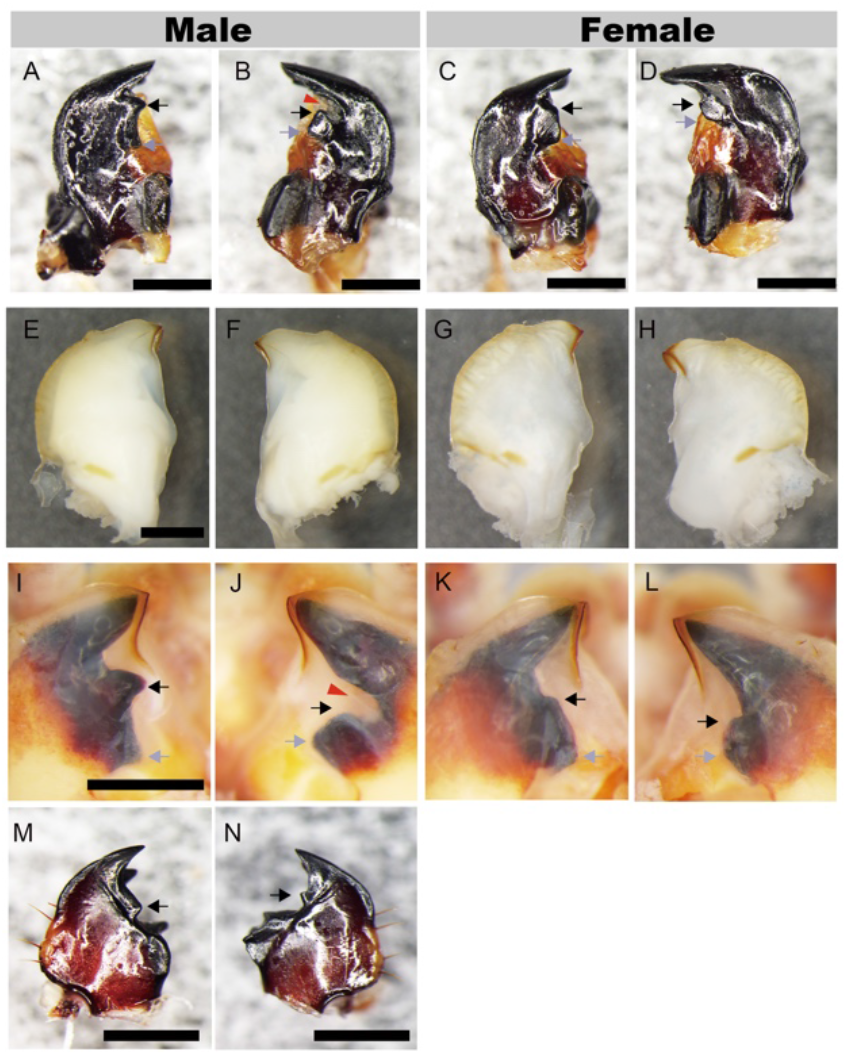
Mandibular morphology. The mandibles of adults (left: A and C, right: B and D), pupae (left: E, G, I, and K, right: F, H, J, and L), and larvae (M and N) are shown. E-H and I-L show day 4 and day 11 pupae, respectively. M and N show the mandibles of old larvae (1.2 g weight). Gray and black arrows indicate the first and second inner teeth, respectively. The red arrowhead indicates the right-specific invaginations of males. The pigmented mandibles of pharate adults within pupal cuticle are shown in I-L. The inner teeth of larval mandibles are shown by arrows in M and N. Larval sex was not discriminated because available morphological characteristics for sex determination have not yet been identified in this species. Scale bars indicate 1 mm.

## MATERIALS and METHODS

### Beetles

The metamorphosis of *Z. atratus* largely depends on larval density, and larval weight also affects metamorphosis in this species (Quennedey et al., 1995, Kim et al., 2015). To efficiently induce metamorphosis, we selected large larvae that weighed between 1100 and 1400 mg, and isolated them into plastic cases (width 4.0 x length 4.5 x height 3.5 cm) with bran as food. Isolated larvae were reared at 25°C, and larval development was observed every day. In *Z. atratus*, isolated larvae take the crooked posture before metamorphosis. It is considered as the prepupal stage. Therefore, this crooked posture was used as the indicator of initiation of metamorphosis.

Pupae (0-5 days after pupal molting) were fixed using 4% paraformaldehyde in PBS, and preserved in methanol until observations of the mandibles. Pupal heads were dissected from the bodies and incubated in 6% hydrogen peroxide in methanol for 12 h for bleaching. The mandibles of pharate adults within the pupal cuticle were drawn out in methanol. Adults and larvae were preserved at -20°C until observations of the mandibles.

*T. castaneum* was reared as described previously (Minakuchi et al., 2008). The wild-type strain of *T. castaneum* was provided by the National Agriculture and Food Research Organization (Tsukuba, Japan). *T. castaneum* was reared at 30°C in the constant dark in a LP-130P incubator (NK system, Osaka, Japan) and fed whole wheat flour. The whole wheat flour including *T. castaneum* larvae were sieved using a 710-μm sieve, and then the larvae remaining on the sieve were picked up as “old larvae”.

### RNA sequencing

Total RNA was extracted using the SV Total RNA Isolation kit (Promega, Madison, WI, USA). Total RNA extracted from the male mandibles on each day (day 0-5 pupa) was mixed for the left and right mandibles separately. We prepared biological duplicates for the left and right mandibles. The quality of RNA was checked using a 2100 Bioanalyzer (Thermo Fisher Scientific, Waltham, MA, USA). cDNA libraries were constructed using TruSeq RNA Sample Prep Kit v2 (Illumina, San Diego, CA, USA). Libraries were prepared by the random fragmentation of cDNA samples, and 5’ and 3’ adapter ligation was then performed. Adapter-ligated fragments were PCR amplified and gel purified. Sequencing was performed using Hiseq 2500 (Illumina) by Macrogen Corp. Japan.

### RNA sequence data analysis

Sequenced RNA-seq read data were deposited into the Data Bank of Japan (DDBJ) SRA. The assigned accession numbers of read data from left mandible biological replicate 1, right mandible biological replicate 1, left mandible biological replicate 2, and right mandible biological replicate 2 were DRR173275, DRR173276, DRR173277, and DRR173275, respectively. RNA-seq reads were cleaned by Trimmomatic-0.36 (Bolger, Lohse, & Usadel, 2014). In the de novo assembly to construct contigs, all of the cleaned sequence reads were used with Trinity-version2.4.0 (Grabherr et al., 2011). Assembled contig data were deposited into the Transcriptome Shotgun Assembly (TSA) database. The accession numbers of the contigs were ICLG01000001-ICLG01091161 (91161 entries). The methods of annotating each contig were previously described by Uchibori-Asano et al. (2019). Briefly, each transcript sequence used as a query against the National Center for Biotechnology Information non-redundant (NCBI-nr) protein database with the blastx method (e-value < 1e-3) and top-hit description was extracted. The candidate coding sequence (CDS) of each contig was predicted with Transdecoder bundled with Trinity r20140717. CDS amino acid sequences were analyzed with the Pfam domain database by HMMER3 (Eddy, 2011).

To estimate the expression abundance of each contig, the “align_and_estimate_abundance.pl” program bundled with Trinity r20140717 using RSEM for the estimating method was used (Haas et al., 2013). The expression abundance data of each sample was deposited into the DDBJ Genomic Expression Archive (Accession number E-GEAD-310). To detect differentially expressed genes (DEGs) between left and right mandible samples, iDEGES/edgeR in the TCC package version 1.8.2 was used with a false discovery rate (FDR) < 0.05 and fold change of normalized tag counts > 2 (Sun et al., 2013).

### RNAi

DNA templates for double-stranded (ds) RNA synthesis were prepared by PCR with primers conjugated with T7 promoter sequences at the 5’ end. The *malE* gene was used as a negative control (Minakuchi et al., 2008). Primers used in this study are listed in Supplementary Table S1. PCR products were purified using the QIAquick gel extraction kit (Qiagen, Venlo, the Netherlands). dsRNA was transcribed by T7 RNA polymerase in the Megascript T7 Kit (Ambion, Austin, TX, USA). dsRNA solution was treated with DNase for 15 minutes and then purified by isopropanol precipitation. dsRNAs (*T. castaneum*: 750 ng, *Z. atratus*: 10 μg) were injected into the boundary between the sternum using the injection holder HI-7 (Narishige, Tokyo, Japan) connected with a glass needle prepared with GD-1 (Narishige). Morphological analysis was carried out in the adult stage. The dsRNA was injected into larvae or day 2 pupae. Larvae injected with dsRNA were reared in the plastic box (Beetles in Materials and Methods). Old larvae (Beetles in Materials and Methods) of *T. castaneum* injected with dsRNA were reared individually in 24-well plates until adult emergence.

Total RNA was extracted from the left and right mandibles of *Z. atratus* pupae using the SV total RNA isolation system (Promega). Extracted RNA was treated with RNase-free DNase I in the SV total RNA isolation system to remove genomic DNA. The quality and concentration of RNA were confirmed using the NanoVue spectrophotometer (GE Healthcare, Chicago, IL, USA). cDNA was synthesized from total RNA (400 ng) with an oligo (dt) primer using the SuperScript III First-strand Synthesis System for RT-PCR (Invitrogen). Gene-specific primers were designed using Primer 3 Plus (Table S1). The suitability of three reference genes [*ribosomal protein L32* (accession no. BBM96696.1), *Elongation Factor1-alfa* (*EF1-α*, accession no. BBM96695.1), and *glyceraldehyde-3-phosphate dehydrogenase* (*GAPDH* (accession no. BBM96697.1)] were evaluated with the software Normfinder (Andersen et al. 2004). Real-time quantitative RT-PCR was performed with PikoReal 96 (Thermo Fisher Scientific, Waltham, MA, USA). Relative expression was calculated by adopting the standard curve method. Expression levels for knockdown verification were calculated from biological replicates (n=3). Statistical analyses were performed using the Student’s *t*-test or Tukey’s test (Statcel, OMS publishing, Tokorozawa, Japan).

### Phylogenetic analysis

Phylogenetic analysis for the homology verification of target genes was performed by MAFFT and PhyML with default settings in NGPhylogeny.fr (Lemoine et al 2019). Phylogenetic trees were visualized by Figtree v1.4.4 (https://github.com/rambaut/figtree accessed by 13 Oct 2023).

### Morphometric analysis for assessment of RNAi effect

Images were taken using the DP22 CCD camera (Olympus, Tokyo, Japan) attached to the SZX7 binocular microscope (Olympus). RNAi effects for morphology were assessed by geometric morphometrics using Geomorph R packages (Baken et al 2021; Adams et al 2023) (https://rdocumentation.org/packages/geomorph/versions/4.0.6, accessed by 13 Oct 2023).

## RESULTS AND DISCUSSION

### Morphological observations of mandibles

The mandibular morphologies of adults (Fig. 1A-D), pupae (Fig. 1E-L), and larvae (Fig. 1M and N) of *Z. atratus* were observed. Inner tooth asymmetry was more prominent in males than in females (Fig. 1A-D, arrows). The inner teeth were composed of first (Fig. 1A-D, grey arrows) and second (Fig. 1A-D, black arrows) inner teeth. Invaginations between the apical teeth and the second inner teeth were obvious in males (Fig. 1B, red arrowhead). The morphology of whole pupal mandibles appeared to be symmetric (Fig. 1E-H), and the asymmetric inner teeth of adults were formed within pupal cuticle (Fig. 1I-L, arrows). Larval inner teeth were also asymmetric (Fig. 1M and N). The number of tips of the larval inner tooth was one in both mandibles; however, the left inner teeth were broader than the right ones (Fig. 1M and N, arrows). These results showed that the extent of inner tooth asymmetry changed throughout postembryonic development.

Since we found that the adult mandibles were able to observe under the pupal cuticle (Fig. 1I-L, arrows), we observed the development processes of adult male mandibles in the pupal cuticle to reveal timing of inner tootth formation. We found that adult mandibular formation was finished in 5 days after pupal molting (Fig. 2). Left second inner teeth were formed within 3 days (Fig. 2A, C, E, K, and M, arrowheads), whereas all right inner teeth were not yet clearly visible (Fig. 2B, D, F, L, and N). Thus, the developmental timing of left second inner teeth was earlier than those of right mandibles. First inner teeth in left and right mandibles were formed in 4-5 days (Fig. 2G-J, O, and P, green arrowheads). Right characteristic invaginations in males were also formed in 4-5 days (Fig. 2P, black arrowhead).

**Figure 2.**
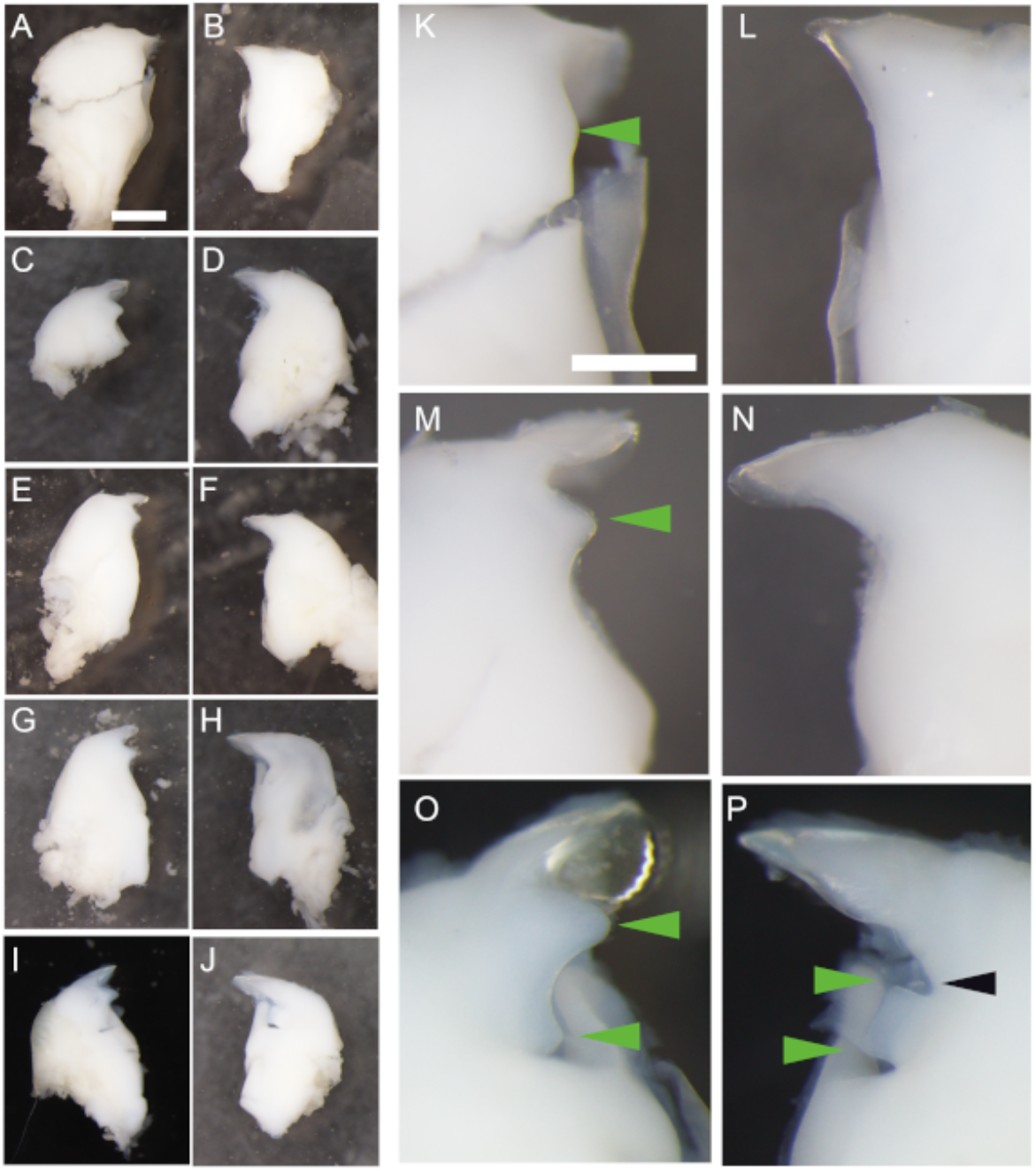
Morphological changes in adult mandibles during male pupal development (day 1 to 5). Mandibles of pharate adults of day 1 (A, B, K, and L), day 2 (C and D), day 3 (E, F, M, and N), day 4 (G and H), and day 5 (I, J, O, and P) pupae were dissected from pupal cuticle. Green arrowheads indicate developing inner teeth. Black arrowheads indicate the right-characteristic invagination. Scale bars show 500 μm.

### Transcriptome in pupal mandibles

Genes with differential expression between the left and right mandibles are thought to be responsible for morphological differences of asymmetric inner teeth. Therefore, we compared gene expression between the left and right mandibles to find these genes. We performed RNA sequencing (RNA-seq) using a mixture of RNA extracted from day 0-5 right or left pupal mandibles (Supplementary Methods). RNA-seq showed no differentially expressed genes (Supplementary Table S2). However, RNA-seq offered sequence information of transcripts that is expressed in the pupal mandible, allowing RNAi analyses of candidate genes as described below.

### The role of candidate genes

We attempted to identify genes involved in the asymmetric formation of the inner teeth by RNAi of potential candidate genes (*paired* (*prd*), *aristaless* (*al*), *LIM homeobox 1* (*Lim1*), and *mille-pattes* (*mlpt*)). In *D. melanogaster, prd* is expressed in the mandibular segment in late embryonic development (Davis et al., 2001). In *T. castaneum, prd* regulates the formation of whole mandibles during embryonic development (Dönitz et al., 2015, Schmitt-Engel et al., 2015). In the stag beetle *C. metallifer, aristaless* (*al*) is involved in inner tooth formation (Gotoh et al., 2017). During leg development of *D. melanogaster, al* expression is affected by *LIM homeobox 1* (*Lim1*) expression (Tsuji et al., 2000). At early germband stage of *T. castaneum*, the expression of *mille-pattes* (*mlpt*) is conspicuous in mandibular segment and posterior end (Savard et al. 2006). These genes are possible to be associated with the inner teeth formation in *Z. atratus*. We identified the homologs of *paird* (ICLG1078576 and ICLG01078581), *aristaless* (ICLG01000824), and *Lim1* (ICLG01004303) by constructing the Maximum likelihood phylogeny. The predicted homologs of *Z. atraus* were assigned into the phylogenetic group that include the corresponding homolog of *D. melanogaster* (Supplementary Figure S1). *mlpt* encodes four short peptides, and the first three and last one contains LDPTGXY and ETSSGRRRR motif (Savard et al., 2006; Tour and McGinnis, 2006). We confirmed the presence of these motif in ICLG01004445 and ICLG01004460, which were defined as a homolog of *mlpt* (Supplementary Figure S2).

We performed RNAi for *prd, al, Lim1, and mlpt* to investigate their functions in the formation of inner teeth (Table 1, Figure 3). The RNAi effects of all these genes on the inner tooth formation were observed with high penetrance (Table 1). RNAi effects of all genes were compared with that of *malE* (Fig. 3A and B) when dsRNAs were injected into larvae. *prd* RNAi resulted in the absence of inner teeth in both the left and right mandibles (Fig. 3C and D). When dsRNA was injected into day 2 pupae, the adult males lacked the characteristic right-mandibular invagination between the apical and second inner teeth, without affecting the morphology of left inner teeth (Fig. 3E and F), indicating that *prd* functions only in the right mandible, depending on the stage of mandibular development. We confirmed the effects of the *prd* knockdown at the gene expression level when we performed larval and pupal injections (Supplementary Figure S3). To reveal the function of *prd* in species possessing different inner tooth morphologies, we performed *prd* RNAi on *T. castaneum*. Adults of this species possess one inner tooth, in contrast to two inner teeth in *Z. atratus* (Supplementary Figure S4A and B, arrows). The knockdown of *prd* of *T. castaneum* resulted in the lack of the one inner tooth in the left and right mandibles (Supplementary Figure S4 C and D). Although it was unable to perform a morphometric analysis because of the too small mandibles of *T. castaneum*, the *prd* knockdown also appeared to affect the apical teeth. These results demonstrated that *prd* was also necessary for the formation of asymmetric inner teeth in *T. castaneum*. RNAi for *al* affected the formation of the first inner teeth in left and right mandibles in *Z. atratus* (Table 1 and Fig. 3G and H, arrowheads). RNAi for *Lim1* not only affected first inner teeth in the same manner as *al* knockdown (Fig. 3I and J, arrowheads), but also strongly affected the second inner teeth, unlike *al* knockdown. *mlpt* RNAi markedly affected the formation of the right second inner teeth (Fig.3 K and L, arrowhead) and apical teeth. RNAi results in females were similar to those in males (Supplementary figure S5).

**Table 1.** RNAi effect of candidate genes on inner teeth formations.

| Gene | The number of animals examined in this study |  | The number of animals in which abnormalities were observed in mandibles by RNAi |  |
| --- | --- | --- | --- | --- |
|  | Male | Female | Male | Female |
| malE | 7 | 4 | 0 | 0 |
| ICLG01078581 (*prd*) | 16 (8) | 5 (5) | 16 (7) | 5 (0) |
| ICLG01000824 (*aristaless*) | 11 | 1 | 11 | 1 |
| ICLG01004303 (*Lim1*) | 3 | 0 | 3 | 0 |
| ICLG01004445, ICLG01004460 (*mlpt*) | 16 | 6 | 11 | 3 |

The numbers in parentheses indicate the number of animals in pupal RNAi experiments.
Footnote: The numbers in parentheses indicate the number of animals in pupal RNAi experiments.

**Figure 3.**
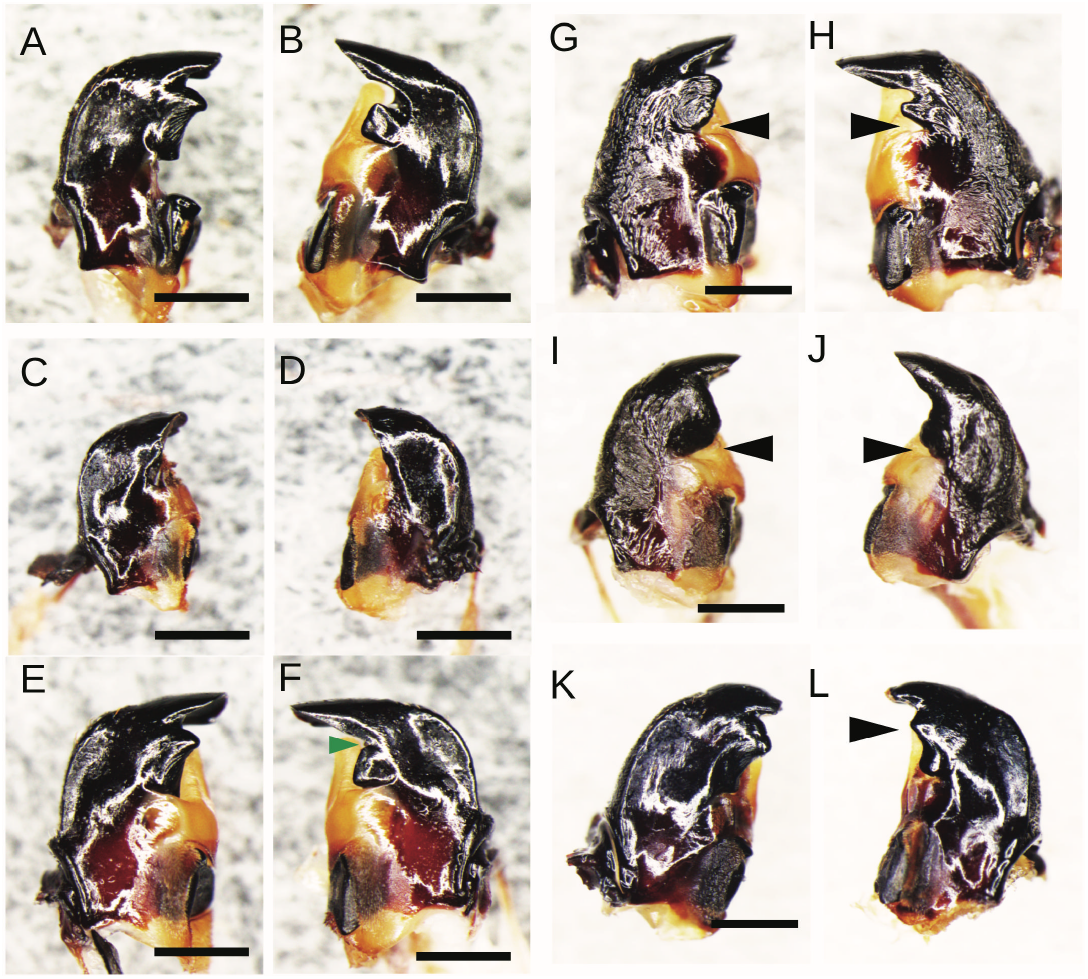
RNAi effects on the mandibular morphologies of male adults. Larval RNAi of *malE* (A, B), larval RNAi of *prd* (C, D), pupal RNAi of *prd* (E, F), larval RNAi of *al* (G, H), larval RNAi of *Lim1* (I, J), and larval RNAi of *mlpt* (K,L). Arrowheads indicate the characteristic morphological regions caused by RNAi of each gene. Scale bars indicate 500 μm.

Geometric morphometrics analysis was performed to explore the morphological parts largely affected by RNAi. The extent of morphological changes in the mandibular base, molar teeth, inner teeth, and apical teeth were quantified, and then PCA was used for the assessment of morphological changes. This analysis was applied to *prd*- and *mlpt*-knockdown adults after dsRNA injection during the larval stages, because their knockdown caused drastic morphological changes in this study (Figure 3). PCA showed that the effect of *prd* RNAi was clearly observed both in left and right mandibles (Figure 4A and 4B). The PC1 values reached 75% both in left and right mandibles. Shape differences between *malE* and *prd* were visualized using maximum and minimum values of PC1, indicating that the morphology of the inner teeth was largely changed In the left and right mandible, while basal regions of mandibles were not changed (Figures 4C and D).

**Figure 4.**
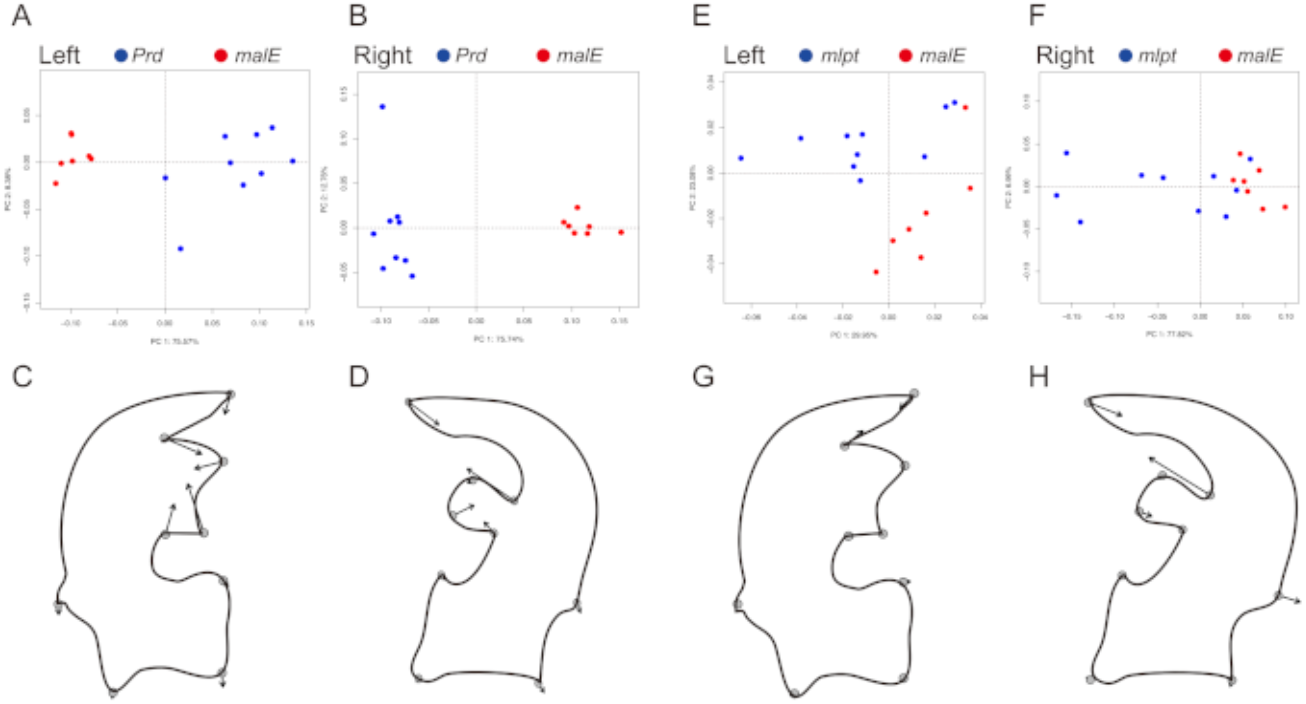
The assessment of morphological changes caused by RNAi. PCA results of *prd* and *mlpt* RNAi (A-D). The plots of each gene is indicated by different colors (A-D). At the bottom of the PCA results, the illustrations of mandibles are shown together with the measurement points. The direction and degree of morphological change induced by RNAi were shown by arrows visualized using the max and minimum values of PC1.

Although not as clearly grouped as the plot of the *prd* knock-down, PCA showed that the *mlpt* RNAi affected the morphologies of several individuals (Figure 4E and F). Although the tips and bases of apical teeth were affected by *mlpt* RNAi in the left mandibles (Figure 4G), the contribution rate of PC1 was low (29.95%). This suggests that while *mlpt* RNAi cause morphological changes in these regions, these changes were not drastic, as a higher PC1 contribution would be expected for more significant morphological alterations. In the right mandibles, large morphological changes due to RNAi were observed in invagination and apical teeth (Figure 4H). High contribution ratio of PC1 (77.82%) shows that *mlpt* RNAi affected the morphology of invagination and apical inner teeth mainly. In particular, the morphological changes at the invagination are remarkable, and that is consistent with the morphological changes of inner teeth seen in Figure 3L. In summary, *prd* is implicated in formation of the inner teeth, and *mlpt* is particularly implicated in the formation of invagination in the right mandibles.

In this study, we aimed to identify genes responsible for morphological differences of asymmetric inner teeth. Unlike results of *prd* gene, *mlpt* RNAi had a strong effect on the morphology particularly in right-specific invaginations, suggesting its involvement in the formation of mandibular asymmetry. RNAi showed that *mlpt* was implicated to form right-specific invaginations locally. This suggest that *mlpt* is the gene only functioning in the right side. In *Tribolium castaneum, mlpt* is responsible for the abdominal segment identity, and *mlpt* knockdown leads to homeotic transformation from abdominal segment to thoracic segment (Savard et al 2006). This is the first report that *mlpt* functions in the formation of the mandible. Several small ORF regions are included in *mlpt* mRNA. In recent years, there is growing evidence that peptides translated from smORFs can be key factors in development and physiology in a variety of organisms (Aspden et al 2014, Plaza et al 2017). Morphogenetic genes show various spatiotemporal expression patterns in embryos, leading to morphological differences along the anterior-posterior, dorsal-ventral, and proximal-distal axes. However, genes that induce asymmetric differences between the left and right sides have not yet been identified. This study showed that *mlpt* may be key regulators generating the asymmetry in mandibles.

### Limitations of this study

Morphological observation showed that inner tooth formation occurs gradually and locally during pupal development, resulting in the asymmetric mandibles. RNAi experiments showed that four genes identified in this study were implicated in the formation of inner teeth. These results suggest that stepwise and locally regulated expression of these four genes is responsible for the formation of the gradually and locally formation of inner teeth. Although we attempted to identify the expression regions of these genes using whole mount in situ hybridization, it was not successful. The specific spatial and temporal expression patterns of these genes during asymmetric inner tooth formation remain to be elucidated for further understanding of the exact molecular mechanisms underlying this process.

## ACKNOWLEDGMENTS

We thank Yuri Homma (Nihon University) for help with laboratory work. This study was supported by personal research expenses of Nihon University.

## DATA AVAILABILITY

Sequenced RNA-seq read data is available in the Data Bank of Japan (DDBJ) SRA (DRR173275, DRR173276, DRR173277, and DRR173278). Assembled contig data is deposited into the Transcriptome Shotgun Assembly (TSA) database. The accession numbers of the contigs were ICLG01000001-ICLG01091161 (91161 entries). Supplementary figures and tables are available at figshare (https://doi.org/10.6084/m9.figshare.26860714.v1).

## AUTHOR CONTRIBUTIONS

Methodology: K.T., K.Y., T.T.; Validation: T.K., Formal analysis: K.T..; Investigation: K.T..; Resources: K.T., T.T; Data curation: K.T., K.T.; Writing - original draft: K.T., K.Y., T.T.; Writing - review & editing: K.T., K.Y., T.T.; Visualization: K.T.; Supervision: K.T.; Project administration: K.T.; Funding acquisition: K.T., T.T.

